# Atypical intrinsic neural timescales in temporal lobe epilepsy

**DOI:** 10.1101/2022.07.01.498416

**Authors:** Ke Xie, Jessica Royer, Sara Lariviere, Raul Rodriguez-Cruces, Reinder Vos de Wael, Bo-yong Park, Hans Auer, Shahin Tavakol, Jordan DeKraker, Chifaou Abdallah, Lorenzo Caciagli, Dani S. Bassett, Andrea Bernasconi, Neda Bernasconi, Birgit Frauscher, Luis Concha, Boris C. Bernhardt

## Abstract

**Objective:** Temporal lobe epilepsy (TLE) is the most common drug-resistant epilepsy in adults. Here, we aimed to profile local neural function in TLE in *vivo*, building on prior evidence that has identified widespread structural alterations. Using multimodal MRI, we mapped intrinsic neural timescales (INT) at rest, examined associations to TLE-related structural compromise, and evaluated the clinical utility of INT.

**Methods:** We studied 46 TLE patients and 44 healthy controls from two independent sites, and mapped INT changes in patients relative to controls across hippocampal, subcortical, and neocortical regions. We examined region-specific associations to structural alterations and explored effects of age and epilepsy duration. A supervised machine learning paradigm assessed utility of INT for classifying patients-*vs*-controls and seizure focus lateralization.

**Results:** Relative to controls, TLE showed marked INT reductions across multiple regions bilaterally, indexing faster changing resting activity, with strongest effects in ipsilateral medial and lateral temporal regions, and sensorimotor cortices. Findings were consistent in each site and robust, albeit with reduced effect sizes, when correcting for structural alterations. TLE-related INT reductions increased with advancing disease duration, yet findings differed from aging effects seen in controls. Classifiers based on INT distinguished patients-*vs*-controls (balanced accuracy, 5-fold: 76±2.65%; cross-site, 72-83%) and lateralized the focus in TLE (balanced accuracy, 5-fold: 96±2.10%; cross-site, 95-97%) with high accuracies and generalization.

**Conclusions:** Our findings robustly demonstrate atypical macroscale function in TLE in a topography that extends beyond mesiotemporal epicenters. INT measurements can assist in TLE diagnosis, seizure focus lateralization, and monitoring of disease progression, which suggests clinical utility.

## Introduction

Temporal lobe epilepsy (TLE) is a common drug-resistant epilepsy in adults. Traditionally considered as a ‘focal’ epilepsy with hippocampal pathology as its hallmark^1^, mounting histological and neuroimaging work indicates large-scale reorganization of brain structure. Structural and diffusion magnetic resonance imaging (MRI) studies, in particular, have revealed gray and white matter alterations across multiple cortical and subcortical areas^2, 3^, supporting the notion that TLE is a network disorder^4, 5^.

Beyond the identified structural alterations, TLE has long been recognized to impact brain function^6, 7^. Resting-state functional MRI (rs-fMRI) is increasingly used to interrogate neural function in health and disease, in light of its high spatial-resolution and full-brain coverage. Prior rs-fMRI studies in TLE identified atypical functional connectivity relative to controls, with altered inter-regional communication within circuits of temporal cortices, as well as between temporal and extra-temporal regions^8-10^. These findings are complemented by connectome-level investigations showing whole-brain functional reorganization^11, 12^. In the literature focusing on functional imbalances in TLE, however, notably few studies have characterized intrinsic function at a regional level. Notably, and beyond the spatial localization of atypical function and connectivity, changes in functional signaling can also be characterized across temporal scales, a marker sensitive to neural processing^13^. One marker increasingly used in prior rs-fMRI and electrophysiology studies in healthy individuals is the measurement of intrinsic neural timescales (INT), which are sensitive to the shape of the long-range temporal autocorrelation of physiological signals^14, 15^. As such, it helps to differentiate faster and slower processing. Moreover, studies have emphasized that INT vary topographically and recapitulate functional zones, with heteromodal association cortices implicated in higher order signal integration longer timescales than sensory/motor and unimodal cortices^14-16^. Notably, an increasing body of studies conducted in non-human primates as well as healthy humans has furthermore shown systematic associations between INT and region-to-region variations in brain morphology and microarchitecture^15, 17^, suggesting that INT could serve as a local functional index anchored in foundational models of brain microcircuit organization. As imbalances in connectivity as well as microcircuit organization are recognized in TLE^10, 18^, the systematic mapping of INT changes at a regional level may inform the development of patient-specific markers of atypical brain function.

Here, we mapped the topography of INT in TLE patients and examined alterations relative to healthy individuals. Aggregating data across two independent sites, we opted for a multimodal MRI acquisition and data processing paradigm data that allowed us to profile the topography of INT changes in TLE-*vs*-controls along hippocampal, neocortical, and subcortical regions. Moreover, integration of the rs-fMRI measures with structural and diffusion MRI data allowed us to explore whether INT alterations in patients relate to brain structure and fiber microstructure in TLE^10, 19^. We also explored associations between INT and age and epilepsy duration, to provide a functional perspective on the notion that TLE may be associated with atypical aging and cumulative disease effects. Finally, we assessed whether brain-wide INT parametrization can inform patient-vs-control classification and seizure focus lateralization, leveraging a supervised learning paradigm that implicated both cross-validation and cross-site generalizability assessments.

## Methods

### Participants

We studied people with TLE and healthy individuals who were aggregated from two independent sites: *a)* the MICA-MICs dataset from the Montréal Neurological Institute and Hospital^20^, and *b)* the EpiC dataset from the Universidad Nacional Autónoma de México^21^.

#### (a) MICA-MICs

There were 19 people with drug-resistant unilateral TLE (8 males; mean±SD age = 37.1±11.5 years, 15/4 left/right-TLE) and 23 healthy individuals (12 males; 34.5±2.9 years). There were no differences in age and sex between both cohorts (**Table 1**; age, *t* = -1.01, *p* = 0.32; sex, *χ*^2^ = 0.42, *p* = 0.52). Diagnosis of TLE and lateralization of seizure foci were determined by the criteria of the International League Against Epilepsy (ILAE), based on a comprehensive examination that included detailed clinical history, neurological examination, review of medical records, video-electroencephalography (EEG) recordings of ictal and interictal events, and clinical MRI evaluation. Age at seizure onset was 22.0±9.7 years and duration of epilepsy was 15.1±12.9 years.

**Table 1.** Demographic and clinical characteristics

|  | *MICA-MICs* |  |  | *EpiC* |  |  |
| --- | --- | --- | --- | --- | --- | --- |
|  | Controls | TLE | *p*-value | Controls | TLE | *p*-value |
| Male/female | 12/11 | 8/11 | 0.52<sup>a</sup> | 8/13 | 10/17 | 0.94<sup>a</sup> |
| Age | 34.5±2.9 | 37.1±11.5 | 0.32<sup>b</sup> | 29.8±11.7 | 29.9±11.5 | 0.98<sup>b</sup> |
| Seizure focus, L/R | - | 15/4 | - | - | 16/11 | - |
| Age at seizure onset | - | 22.0±9.7 | - | - | 13.4±9.4 | - |
| Disease duration | - | 15.1±12.9 | - | - | 16.5±13.3 | - |

Age, age at seizure onset, and disease duration are presented in mean±SD years. TLE, temporal lobe epilepsy; L, left; R, right.
Note: <sup>a</sup>, Chi-square test; <sup>b</sup>, Two-sample *t*-test.

#### (b) EpiC

There were 27 people with drug-resistant unilateral TLE (10 males; mean±SD age = 29.9±11.5 years, 16/11 left/right-TLE) and 21 age-and sex-matched healthy controls (8 males; 29.8±11.7 years). There were no differences in age and sex between both cohorts (**Table 1**; age, *t* = -0.03, *p* = 0.98; sex, *χ*^2^ = 0.01, *p* = 0.94). Patients were diagnosed by certified neurologists based on ILAE criteria as detailed above. Age at seizure onset was 13.4±9.4 years and duration of epilepsy was 16.5±13.3 years.

Considering the TLE patients, we observed no between-site differences with respect to sex (*χ*^2^ = 0.12, *p* = 0.73), the proportion of patients with a left-/right-sided seizure focus (*χ*^2^ = 1.97, *p* = 0.16), and epilepsy duration (*t* = 0.35; *p* = 0.73). On the other hand, there were differences in age (*t* = -2.05, *p* = 0.05) and age at seizure onset (*t* = -2.95, *p* = 0.005).

### Standard protocol approvals, registrations, and patient consents

The studies had been approved by the Research Ethics Boards of the Montreal Neurological Institute and Hospital and of the Universidad Nacional Autónoma de México, respectively. Written informed consent was obtained from all participants in accordance with the Declarations of Helsinki.

### MRI acquisition

#### (a) MICA-MICs

Data were collected on a 3T Siemens Magnetom Prisma-Fit equipped with a 64-channel head coil. The aquisition included (i) one or two T1-weighted (T1w) scans (3D-magnetization-prepared rapid gradient-echo (MPRAGE) sequence, voxel size = 0.8mm^3^, matrix = 320×320, repetition time (TR) = 2300ms, echo-time (TE) = 3.14ms, flip angle = 9°, inversion time (TI) = 900ms, iPAT = 224 slices), (ii) one rs-fMRI scan (multiband accelerated 2D-BOLD echo-planar imaging (EPI) sequence, voxel size = 3mm^3^, TR = 600ms, TE = 30ms, flip angle = 52°, field of view (FOV) = 240×240mm^2^, multi-band factor = 6, echo spacing = 0.54ms, 48 slices, 700 volumes), and (iii) one diffusion-weighted imaging (DWI) scan (2D spin-echo EPI, voxel size = 1.6×1.6×1.6mm^3^, TR = 3500ms, TE = 64.40ms, flip angle = 90°, FOV = 224×224mm^2^, echo spacing = 0.54ms, 3 b0 images). The DWI consisted of 3 shells with b-values 300, 700, and 2000 s/mm^2^, and 10, 40, and 90 diffusion directions per shell, respectively. During the rs-fMRI scan, participants were instructed to lie still, fixate a cross presented on the screen, and not to fall asleep.

#### (b) Epic

Data were acquired on a 3T Philips Achieva TX scanner equipped with a 32-channel head coil. The acquisition included (i) one T1w scan (3D spoiled gradient-echo, voxel size = 1mm^3^, TR = 8.1ms, TE = 3.7ms, FOV = 179×256×256mm^3^, flip angle = 8°, 240 slices), (ii) one rs-fMRI scan (gradient-echo EPI, voxel size = 2×2×3mm^3^, TR = 2000ms, TE = 30ms, flip angle = 90°, 34 slices, 200 volumes), and (iii) a DWI scan (2D EPI, voxel size = 2×2×2mm^3^, TR = 11.86s, TE = 64.3ms, FOV = 256×256×100mm^3^, b-value = 2000 s/mm^2^, 60 diffusion directions, 2 b0 images). Participants were instructed to keep their eyes closed, stay still and relaxed, and not to fall asleep.

### Multimodal MRI preprocessing

Processing was based on *micapipe* (http://micapipe.readthedocs.io), an open-access multimodal MRI processing pipeline that incorporates structural, rs-fMRI, and diffusion MRI processing streams and allows for cross-modal feature fusion^22^. In brief, *micapipe* automatically segmented subcortical and cortical regions using FreeSurfer (https://surfer.nmr.mgh.havard.edu/)^23^ and FSL-FIRST^24^ on T1w MRI. As in prior studies^25, 26^, cortical thickness was measured as the Euclidean distance of corresponding vertices on pial and white matter surfaces. The hippocampus was segmented using HippUnfold (https://hippunfold.readthedocs.io/)^27^, an open-access hippocampal unfolding and subfield segmentation method that combines a U-Net deep convolutional neural network and topologically-constrained subfield labelling. DWI data were processed using MRtrix (http://www.mrtrix.org/)^28^, incorporating denoising and correction for susceptibility distortions, head motion and eddy currents, followed by fractional anisotropy (FA) and mean diffusivity (MD) estimation. To estimate superficial white matter (SWM) microstructure, we automatically generated a surface 2mm below the grey-white matter boundary^10, 19^, mapped it to DWI space using a boundary-based registration, and sampled FA and MD values. Resting-state fMRI data were processed using FSL (https://fsl.fmrib.ox.ac.uk/fsl/fslwiki/)^24^ and AFNI (https://afni.nimh.nih.gov/afni)^29^. Preprocessing included discarding the first five volumes, reorientation, skull stripping, motion and distortion correction, removal of nuisance signals and artefacts using an ICA-FIX classifier, and regression of automatically identified, high-motion timepoints. A boundary-based registration mapped functional timeseries to each participant’s cortical surface^30^. In parallel, subject-specific subcortical and hippocampal segmentations were non-linearly registered to each participant’s native fMRI space. Functional data were mapped to participant-specific cortical surfaces, then registered to the hemisphere-matched Conte69 surface template, and smoothed using a 10mm full-width-at-half-maximum kernel. Smoothed data were downsampled to a template with 10,000 vertices for computational efficiency. Hippocampal and subcortical timeseries were averaged within the corresponding co-registered segmentations.

### Intrinsic neural timescale maps

We estimated INT values at each cortical vertex and in each subcortical region and the hippocampus following prior work^31, 32^. The autocorrelation function (ACF) of the rs-fMRI timeseries for a given measurement point was calculated using the following formula:

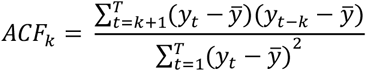

For a given region, *y* denotes the rs-fMRI signal,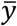 is the mean signal across timepoints, *y* is the time lag (time bin = TR), and *T* is the number of timepoints. Then, INT was calculated as the sum of *ACF* values in the initial positive period:

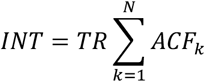

where *TR* is the repetition time of the fMRI signal and *N* is the lag directly preceding the first negative ACF value. Here, multiplying the obtained sum of *ACF* values by *TR* aimed to adjust for differences in the temporal resolution of the rs-fMRI signal. The procedure was repeated for all vertices, subcortical structures and bilateral hippocampi, yielding a whole-brain INT map. As for main analyses, INT maps in patients were *z*-normalized relative to controls across two sites (MICA-MICs & EpiC) and sorted into ipsilateral/contralateral to the seizure focus.

### Statistical analysis

#### i) Case-control analysis

Statistical analysis was carried out using SurfStat (https://mica-mni.github.io/surfstat/) ^33^ for MATLAB [R2021b, The Mathworks Inc.]. As in previous work^10, 25^, we fitted surface-based linear models to compare INT values between TLE and controls, controlling for age, sex, and site:

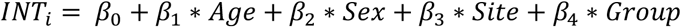

For a given region *i, INT*_*i*_ is the INT measure, *Age* and *Sex* are terms controlling for age and sex, respectively. *Site* is the term controlling for site (*i*.*e*., EpiC and MICA-MICs) and *Group* is the group factor (*i*.*e*., TLE and controls). We also separately evaluated consistency of findings within each site, repeating the above group comparisons while controlling for age and sex:

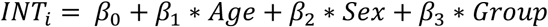

#### ii) Control for cortical morphology and microstructure

To assess INT alterations in TLE patients independent from grey matter morphological and SWM microstructural changes, we examined case-control INT differences while additionally correcting for cortical thickness and SWM diffusion features (*i*.*e*., FA, MD) at each vertex separately^10, 19^. For subcortical structures and the hippocampus, we controlled for the corresponding volume and average diffusion parameters in the surrounding shell^18, 25^.

#### iii) Age-and disease duration-related effects

To study interactions between diagnostic group and age on INT, we first built linear models that included an *age* and *group* main effect term, and an *age×group* interaction term. We also separately evaluated effects of age on INT within TLE and controls. To infer cumulative disease factors in TLE, we assessed effects of disease duration and age of seizure onset on INT.

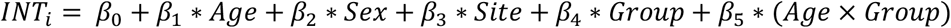

#### iv) Sensitivity analyses

We performed several sensitivity analyses to demonstrate robustness and consistency of our main findings. First, we repeated the INT analysis while controlling for head motion, after calculating mean framewise displacement (FD) of each participant during the rs-fMRI scan^34^. We also repeated the INT analysis while additionally regressing out the global mean signal.

#### v) Correction for multiple comparisons

Findings were corrected for multiple comparisons by controlling the family-wise error (FWE) at *p*_FWE_ < 0.05 for the cortex using random field theory and false discovery rate (FDR) at *p*_FDR_ < 0.05 for subcortical structures and the hippocampus, respectively.

### Spatial permutation tests

Statistical significance of spatial correlations between cortical maps was assessed using spin permutation tests that accounted for spatial autocorrelation^35^, implemented in BrainSpace (https://brainspace.readthedocs.io/)^36^. Specifically, spin tests generated null models (1,000 repetitions) of brain maps by randomly rotating the maps and reassigning each vertex with the values of its nearest neighbors^35^. The significance of the real correlation coefficient was estimated as its position in the null distributions. A similar framework assessed significance of spatial correlations between subcortical maps with the exception that subcortical labels were randomly shuffled while preserving the pairwise three-dimensional Euclidean distance, using the ENIGMA Toolbox (https://enigma-toolbox.readthedocs.io/)^37^.

### Classification and lateralization analysis

To explore whether INT could discriminate patients from controls and lateralize the seizure focus in patients, we utilized support vector machines implemented in LIBSVM (https://www.csie.ntu.edu.tw/~cjlin/libsvm/)^38^. To reduce overfitting, we parcellated cortex-wide INT maps using the Schafer-300 atlas^39^, and subsequently generated inter-hemispheric asymmetry maps, computed as *AI* = *ipsi* - *contra*^40^, where *ipsi* and *contra* was the INT of ipsilateral and contralateral areas. The latter procedure was also applied to subcortical regions and the hippocampus, which yielded 7 additional features. Two models were thus evaluated: *(i)* cortical features only and *(ii)* cortical, subcortical, and hippocampal features. Model performance was quantified in terms of balanced accuracy (BACC) and area under the curve (AUC) of the receiver operating characteristic (ROC). Considering that informative features differed slightly from fold to fold, we obtained feature weights by averaging all folds’ feature weights and then normalized them across parcels (*i*.*e*., |*w*_*i*_|/ ∑ |*w*_*i*_| with *i* indicating *i*^th^ parcel). In the classification analysis, we *z*-normalized asymmetry maps in patients with respect to controls and sorted feature data into ipsilateral/contralateral to the focus, and regressed out age, sex, and site. In the lateralization analysis, we did not *z*-normalize asymmetry maps. Here, we first generated a general linear model for the training set to reduce the effects of age, sex, and site, and applied the estimated parameters to the testing set. The residuals served as algorithm inputs. We assessed classification and lateralization algorithms in two different scenarios. First, we used 5-fold cross-validation with 100 iterations across both datasets combined. Participants across both datasets were randomly split into 5 folds. Classifiers were trained on 4 folds and tested iteratively on the one held-out until all had served as a testing set; this procedure was repeated 100 times so that we can obtain different training and test sets. AUC and balanced accuracy were measured for each repetition and averaged across them. Statistical significance was assessed using 1,000 permutation tests with a threshold of *p*<0.05. Briefly, the features were unchanged, and labels were randomly shuffled for 1,000 times and split into the training and testing sets. The *p* value was calculated by dividing the shuffled times by the number that was equal to or higher than the real balanced accuracy or AUC. Second, we also evaluated cross-site generalizability. Here, we trained the classification/lateralization algorithms on one dataset using leave-one-subject-out cross validation and tested them on the other dataset.

### Code and data availability

Processing pipelines are openly available on https://github.com/MICA-MNI/micapipe and http://micapipe.readthedocs.io. The SurfStat toolbox used for surface-based statistical analysis can be found at http://math.mcgill.ca/keith/surfstat/ or https://mica-mni.github.io/surfstat/. The LIBSVM toolbox can be found at https://www.csie.ntu.edu.tw/~cjlin/libsvm/index.html. Code used in the main analyses are openly available on http://github.com/MICA-MNI/micaopen/. Surface-based INT features are available at https://osf.io/m4fap/.

## Results

### Group differences in INT

We first assessed the topography of INT measures in healthy controls, and observed relatively long INT in parieto-occipital cortices, and shorter INT in sensorimotor and temporo-limbic cortices. In other words, higher-order association areas showed slower processing timescales compared to sensorimotor areas with faster signal fluctuations^14, 15^. As for the subcortical structures, longer INT were seen in the thalamus, and shorter INT in the amygdala, nucleus accumbens and pallidum (**Figure 1A**).

**Figure 1.**
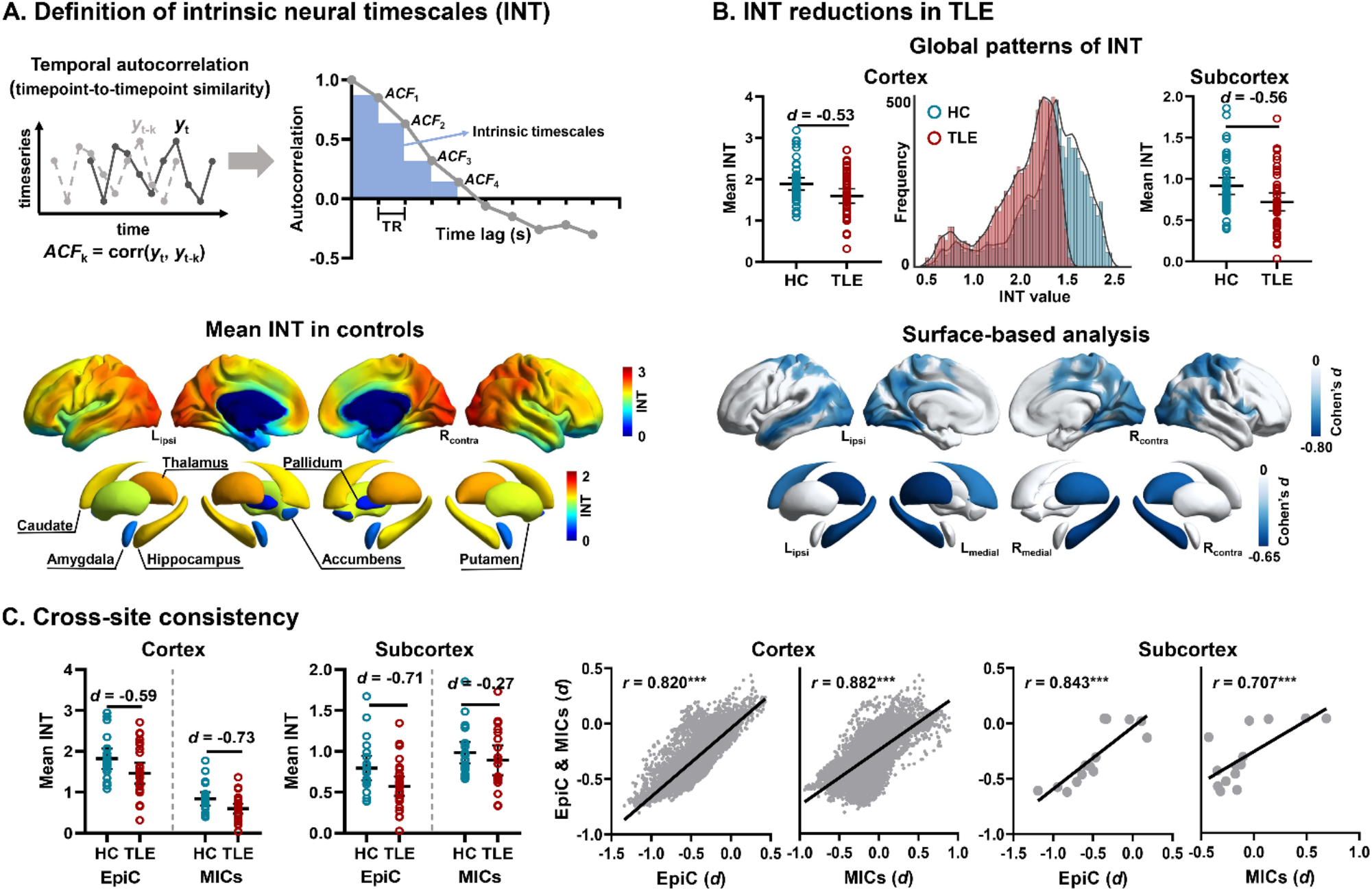
Intrinsic neural timescales (INT) reductions in temporal lobe epilepsy (TLE). **(A)** Calculation of INT from resting-state fMRI data. INT is defined as the sum of timepoint-to-timepoint autocorrelation function (ACF) values in the initial positive period (*i*.*e*., the sum of the area of the blue bars) and multiplied by the repetition time (TR)^31, 32^. Example plots of empirical ACF values for resting-state fMRI data of EpiC site where TR was 2s. Surface-wide maps in the bottom panel show the cortical and subcortical distribution of INT values in healthy controls. **(B)** Differences in INT between TLE patients and controls, controlling for age, sex, and site. *Top*: Global mean and distributions of regional INT. *Bottom*: INT reductions in people with TLE relative to controls after correcting for multiple comparisons at a family-wise error (FWE) rate < 0.05 for cortices, and at a false discovery rate (FDR) < 0.05 for subcortical structures and the hippocampus. **(C)** Cross-site consistency. *Left*: Between-group differences in global mean INT within each site (*i*.*e*., EpiC/MICs). Each circle represents the global mean INT for each participant. *Right*: Correlations of the Cohen’s *d* value from the between group comparison from the overall sample (*i*.*e*., EpiC & MICs combined) and the Cohen’s d from each site separately. Each dot represents an individual vertex or subcortical/hippocampal structure. ***, *p* < 0.001.

Globally, neocortical, subcortical, and hippocampal INT measures were reduced in TLE relative to controls (**Figure 1B**; Cohen’s *d* for neocortex = -0.53; subcortex and hippocampus *d* = -0.56). Local-level analysis found widespread INT reductions in TLE, with highest effects in bilateral precuneus, postcentral and occipital cortices, and ipsilateral middle temporal cortex (*p*_FWE_ < 0.05, Cohen’s *d* = -0.48±0.09), along with bilateral hippocampi and thalamus (**Figure 1B**; *p*_FDR_ < 0.05, Cohen’s *d* = -0.56±0.07). In other words, TLE patients showed faster activity fluctuations in these regions compared to controls.

Findings were relatively consistent across sites, albeit with subtle differences in the magnitude of effect sizes (**Figure 1C** and **Figure S1**; *EpiC:* Cohen’s *d* cortex = -0.59; subcortex and hippocampus = -0.73; *MICA-MICs* = -0.36/-0.27). Effect size measures obtained by pooling data of both sites also significantly correlated with those derived from a single site only (*EpiC:* cortex *r* = 0.82, *p*_spin_ < 0.001; subcortex and hippocampus *r* = 0.84, *p*_shuf_ < 0.001; *MICA-MICs: r* = 0.88/0.71, *p*_spin_/*p*_shuf_ < 0.001).

Our results were robust with respect to head motion and other imaging processing parameters. Specifically, we repeated the INT analysis while controlling for participant’s head motion (*i*.*e*., mean FD during rs-fMRI scans) and observed virtually identical effects (**Figure S2**; cortex: *r* = 0.95, *p*_spin_ < 0.001, *d* = -0.51±0.08; subcortex and hippocampus: *r* = 0.91, *p*_shuf_ < 0.001, *d* = -0.60±0.07). Repeat analyses that additionally regressed out the global rs-fMRI signal, yielded consistent findings for neocortical regions (*r* = 0.76, *p*_spin_ < 0.001; Cohen’s *d* mean±SD = -0.56±0.18) as well as for subcortical regions and the hippocampus (**Figure S2**; *r* = 0.88, *p*_shuf_ = 0.003; Cohen’s *d* mean±SD = -0.46±0.05).

### Effects of cortical morphology and microstructure

We repeated INT analyses after additionally controlling for MRI-based measures of grey matter morphology and SWM microstructure. As for grey matter changes, TLE showed bilateral cortical and subcortical atrophy relative to controls, with strongest effects in paracentral, dorsomedial, and ventromedial prefrontal regions (*p*_FWE_ < 0.05) as well as the ipsilateral hippocampus (*p*_FDR_ < 0.05) (**Figure 2A**), in keeping with prior findings^10, 12, 25^. After controlling for grey matter changes, patterns of INT reductions were similar to our original findings (**Figure 2A**; neocortex: *r* = 0.998, *p*_spin_ < 0.001; subcortex and hippocampus: *r* = 0.933, *p*_shuf_ < 0.001). Effect sizes for INT differences were however reduced both at the neocortical level (Cohen’s *d* = -0.47±0.09, 2% reduction) and at the level of subcortical and hippocampal regions (Cohen’s *d* = -0.47±0.10, 16% reduction). Effect size reductions after controlling for gray matter atrophy were particularly strong in the ipsilateral hippocampus (48% reduction).

**Figure 2.**
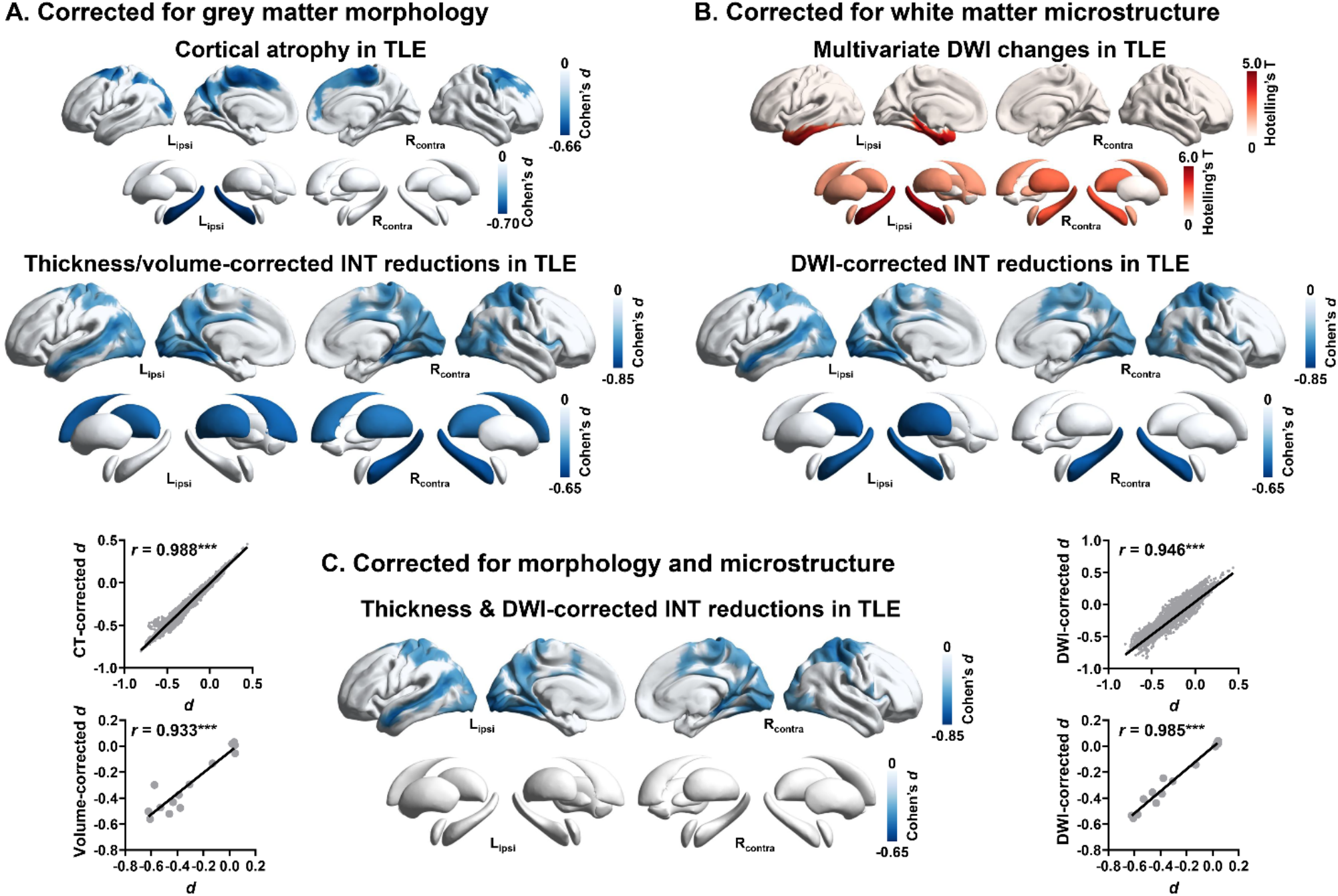
Robustness of intrinsic neural timescales (INT) reductions to grey matter morphological and white matter microstructural alternations. **(A)** Regional cortical thickness and subcortical volume changes in people with TLE compared to healthy controls. INT reductions were similar after controlling for cortical thickness. Similar findings were seen for subcortical structures and the hippocampus. Scatterplots of Cohen’s *d* values from original INT reductions (x-axis; see **Figure 1B**) *vs*. thickness-/volume-corrected INT reductions (y-axis). **(B)** Multivariate superficial white matter changes in TLE relative to controls. INT reductions remained robust when controlling for diffusion effects at each vertex, or surrounding subcortical structures and the hippocampus. Scatterplots of Cohen’s *d* values from original INT reductions (x-axis; see **Figure 1B**) *vs*. DWI-corrected INT reductions (y-axis). **(C)** INT reductions in TLE compared to controls after controlling for both grey matter morphological and white matter microstructural effects. Findings have been corrected for multiple comparisons (See **Figure 1** for details). *, *p* < 0.05; **, *p* < 0.01; ***, *p* < 0.001.

As for SWM diffusivity, multivariate Hotelling’s T tests identified microstructural alternations in TLE, characterized by reduced FA and increased MD (**Figure S4**), in ipsilateral temporo-limbic cortices (*p*_FWE_ < 0.05) and in areas surrounding the bilateral hippocampi and thalamus (*p*_FDR_ < 0.05; **Figure 2B**), also in keeping with prior findings^10, 18, 19^. When correcting for SWM diffusion features, while spatial patterns of INT changes remained similar to our main findings in both neocortex (*r* = 0.95, *p*_spin_ < 0.001) as well as subcortex and hippocampus (*r* = 0.99, *p*_shuf_ < 0.001), effect sizes sightly decreased (Cohen’s *d =* -0.46±0.09/-0.48±0.09). Finally, correcting for both grey matter morphological and SWM microstructural changes did not modify the overall spatial patterns of findings, but reduced effect sizes, particularly in subcortical structures and hippocampus (**Figure 2C**; *r* = 0.93, *p*_shuf_ < 0.001; Cohen’s *d* = -0.45±0.06, 20% reduction) and marginally in the neocortex (*r* = 0.99, *p*_spin_ < 0.001; *d* = 0.45±0.09, 6% reduction).

### Effects of age and disease duration on INT

Globally, we observed significant negative correlations between age and mean cortical and subcortical INT in people with TLE after controlling for sex and site (**Figure 3A**; *r* = -0.333/-0.331, *p* = 0.027/0.025), but not in controls (*r* = 0.116/-0.103, *p* = 0.464/0.505). Moreover, there was a significant discrepancy in cortical slopes between TLE and controls (*z* = -2.120, *p* = 0.017), supporting that the relationship between INT reductions and aging was different in TLE compared to controls. Point-wise interactions between age and group confirmed the above findings, and pointed to clusters of significant interactions in bilateral sensorimotor and occipital regions together with regions in the ipsilateral temporal lobe (**Figure 3A**; *p*_FWE_ < 0.05). *Post hoc* analyses showed that INT increased in these regions in controls with advancing age, while the inverse was observed in TLE (**Figure 3A and 3B**). INT correlated negatively with duration of epilepsy in TLE, with prominent effects in bilateral temporal and prefrontal cortices, and ipsilateral precuneus and central regions (**Figure S5**; *p*_FWE_ < 0.05). There was a trend for negative main effects of disease duration on subcortical INT, particularly in bilateral thalamus (**Figure S5**; *p*_FDR_ < 0.075). In TLE, there were no significant main effects of age at seizure onset on neocortical, subcortical, or hippocampal INT after correction for multiple comparisons (*p*_FDR_/*p*_FWE_ < 0.05).

**Figure 3.**
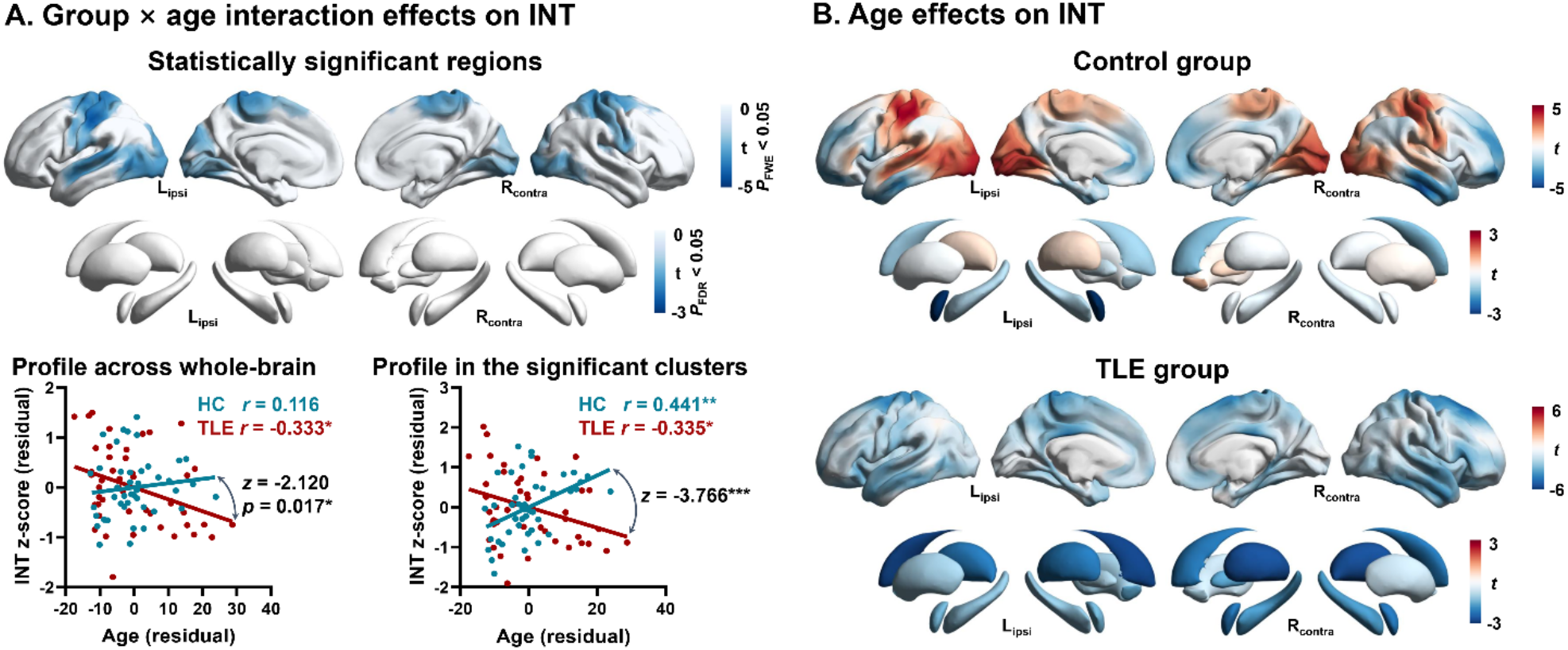
Effects of age on intrinsic neural timescales (INT). **(A)** Interaction between age and group. *Top*: Regions showing a more marked negative effect of age on INT in TLE compared to controls after multiple comparisons correction (See **Figure 1** for details). **(B)** Main effects of age on INT in controls and TLE separately, with INT increasing/decreasing as aging shown in red/blue. *, *p* < 0.05; **, *p* < 0.01.

### Classification and lateralization performance

We leveraged supervised statistical learning to distinguish patients from controls and lateralize the seizure focus in TLE. In the classification analyses, the combination of cortical and subcortical INT achieved a balanced accuracy of 76±2.65% (*p* = 0.001) and AUC of 0.82±0.02 (*p* = 0.001) (**Figure 4A** and **Table 2**). Selected features mapped onto temporo-polar, parahippocampal, medial orbitofrontal, and paracentral regions (**Figure 4A**). In the lateralization analysis, the overall balanced accuracy was 96±2.10% (*p* = 0.001) and the AUC was 0.99±0.01 (*p* = 0.001), based on the combination of cortical and subcortical INT. Informative features were found in the paralimbic cortex, encompassing both temporal pole and insula (**Figure 4B**). Classifiers generalized well from one site to the other (**Figure 4C** and **4D, Table 2)**. Moreover, classifiers based on cortical features alone showed similar performance to that of whole brain models (**Figure S6** and **Table S1**).

**Figure 4.**
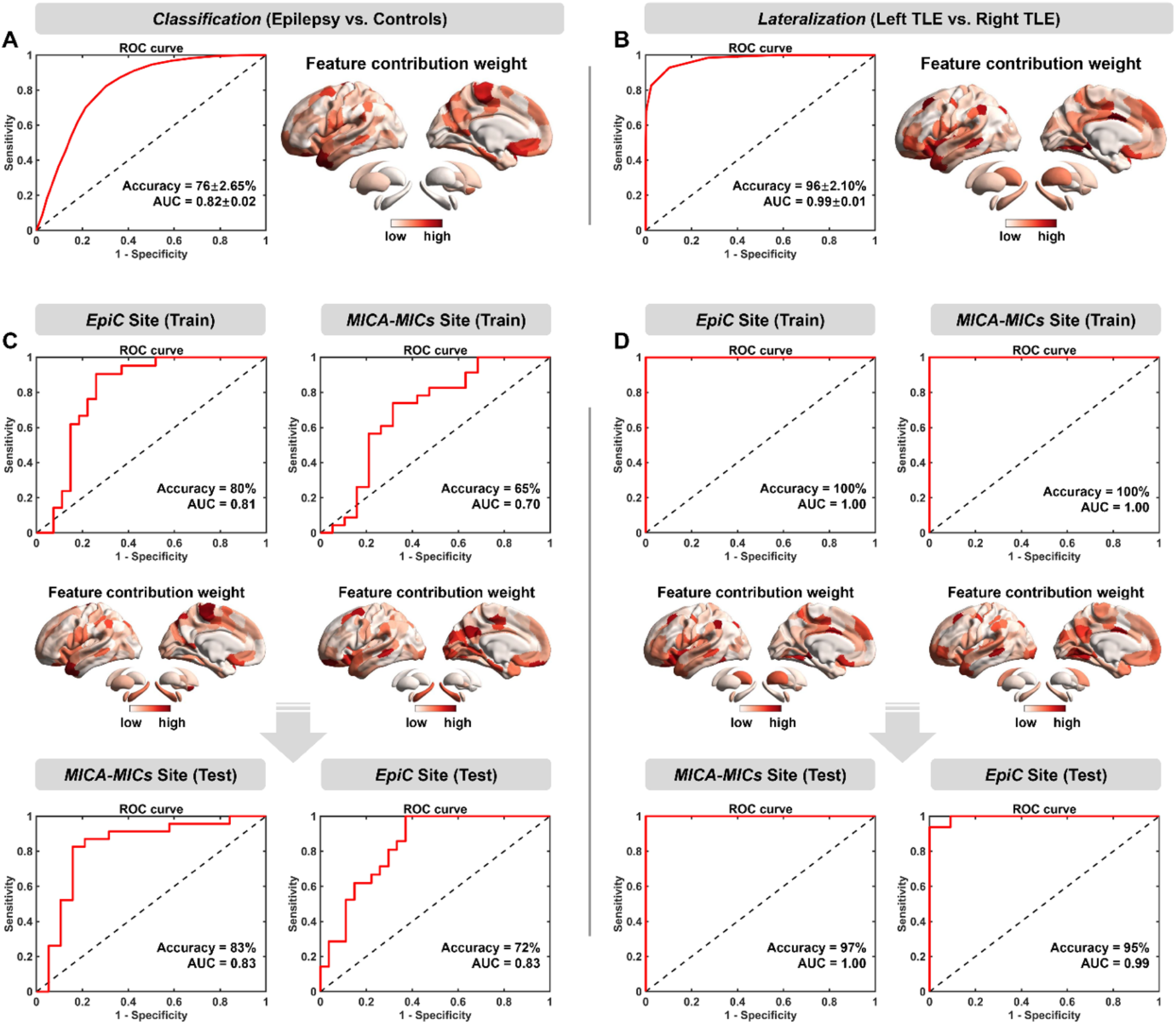
Classification and lateralization analyses using cortical and subcortical intrinsic neural timescales (INT). Support vector machines (SVM) were utilized to classify people with TLE-*vs*-controls (left panel), and to lateralize left-*vs*-right TLE (right panel). **(A)** and **(B)** Performance of classification and lateralization models trained and tested across two datasets using a five-fold cross validation strategy. Average receiver operating characteristic (ROC) curves, normalized feature weight maps, balanced accuracy, and area under the curve (AUC) (mean ± SD) across 100 iterations are indicated. **(C)** and **(D)** Performance of classification and lateralization models trained on one dataset using a leave-one-out cross validation strategy and tested on another one.

**Table 2.** Classification and lateralization performance using combined cortical and subcortical features.

|  | *EpiC & MICs* |  | *EpiC (Train) → MICs (Test)* |  |  |  | *MICs (Train) → EpiC (Test)* |  |  |  |
| --- | --- | --- | --- | --- | --- | --- | --- | --- | --- | --- |
|  | BACC<sup>a</sup> | AUC<sup>a</sup> | BACC | AUC | BACC | AUC | BACC | AUC | BACC | AUC |
| Classification | $76 \pm 2.65\%$ | $0.82 \pm 0.02$ | 80% | 0.81 | 83% | 0.83 | 65% | 0.70 | 72% | 0.83 |
| Lateralization | $96 \pm 2.10\%$ | $0.99 \pm 0.01$ | 100% | 1.00 | 97% | 1.00 | 100% | 1.00 | 95% | 0.99 |

Abbreviations: BACC = balanced accuracy; AUC = area under curve.
<sup>a</sup> BACC and AUC in mean ± SD.

## Discussion

The current work set out to *(i)* map the topography of atypical intrinsic neural timescales (INT) in people with temporal lobe epilepsy (TLE) relative to healthy controls, *(ii)* examine associations to structural alterations in both the grey and superficial white matter, and *(iii)* to assess whether INT changes are sensitive to cumulative disease effects and help in patient-vs-control classification as well as seizure focus lateralization. Studying a two-site dataset of TLE patients and controls, we observed a widespread pattern of INT reductions in TLE, with marked effects in temporo-limbic as well as sensory and motor regions. Results remained consistent, albeit of smaller effect sizes, after controlling for gray matter morphological and white matter microstructural variations in a region-specific manner. Furthermore, correlations analyses with age and disease duration revealed that INT effects aggravated with advanced age and disease duration in patients, a finding supportive of progressive alterations in regional brain function. Finally, supervised classifiers informed by INT features successfully discriminated TLE patients from controls and lateralized the seizure focus with high accuracy and generalization. Collectively, our findings outline a marked and distributed alteration of intrinsic neural function in TLE, which may advance our understanding of structure-function relationships and the development of functional disease biomarkers in the condition.

Core to our work was the quantification of INT from resting-state fMRI (rs-fMRI), a recently developed measure that taps into the temporal autocorrelation of neural signals^14, 15^. In our controls, this measure differentiated sensory and motor systems from heteromodal association cortices, which had longer timescales, in agreement with previous work in healthy adults^14-16^ and developing cohorts^31^. Differences in timescales have previously been related to functional hierarchy of the human cortex^15^. In particular, shorter INT in sensory and motor systems have been associated to their more specialized, externally-oriented functional roles, which may facilitate neural responsiveness to changing environmental contexts. On the other hand, longer INT in heteromodal regions may contribute to their role in multisensory integration as well as abstract, higher-order processing^15, 16^. In typical development, prior studies have shown changes in INT with aging, in particular a lengthening of intrinsic timescales in association and limbic cortices^41^, which may parallel ongoing changes in the microstructural and connectional differentiation in these regions throughout childhood and adolescence^42, 43^. To identify regional patterns of functional imbalances in TLE, we fit neocortical, hippocampal, and subcortical models that additionally controlled for effects of age, sex, and site. This provided robust evidence for marked INT reductions in TLE, in a cortical-subcortical territory encompassing the hippocampus, and thalamus, alongside with lateral temporal, parieto-occipital, and paracentral regions^12^. Effects were overall bilateral, yet more extensive ipsilaterally. This was particularly observed in temporal regions, where INT extended anteriorly to temporopolar areas. In our work, findings of INT reductions were invariably preserved when additionally correcting for head motion and the global signal, suggesting that the results reflected real group differences and not spurious results driven by motion artefacts or imaging processing parameters. Moreover, despite slight differences in effect sizes, patterns of reduced INT in TLE were replicable across independently acquired samples, thus being robust to differences in acquisition protocols and some between site differences in clinical and socio-demographic inclusion.

Prior rs-fMRI studies investigated atypical inter-regional connectivity, and showed atypical functional interactions within mesiotemporal regions^44, 45^, between mesial and lateral temporal regions^7^, and in networks including both temporal and extratemporal regions^9, 44^. In addition, several studies suggest alterations in signal amplitude^19^ or regional signaling homogeneity of neural signals^46^. By robustly demonstrating TLE-related INT reductions across a broad neocortical, hippocampal, and subcortical territory, our work expands this literature by identifying a functional signature characterized by changes in both the spatial and temporal/frequency domain. Temporal lobe structures, in particular the mesiotemporal region as well as the thalamus have long been recognized as critical nodes in the TLE network.^47^ Prior research based on intracranial EEG explorations in human patients together with experimental work in animal models have supported that the hippocampus and adjacent mesiotemporal as well temporopolar regions are highly epileptogenic. Moreover, these regions as well as thalamo-cortical loops play a main role in the propagation of both inter-ictal spiking activity as well as TLE seizures to more distributed cortical networks^18, 48^. In the absence of parallel explorations of electrical activity in the same patients, we cannot confidently say in how far the pattern of INT changes may ultimately predispose brain networks to transient epileptic phenomena. Yet, it is tempting to speculate that the overall reduction in INT across multiple key nodes in the TLE network may reflect a chronic functional imbalance that may more likely generate epileptic phenomena in its dynamic regime. Combining transcriptomics with in vivo functional imaging in healthy individuals, recent work has shown that region-to-region variations in INT align with the expression of genes coding for inhibitory as well as excitatory receptors^15^, which may suggest associations to local excitation and inhibition (E/I) ratio^15, 49^, a finding that is line with computational studies in humans and non-human primates^50^. Our group recently leveraged connectome-informed brain models in TLE and also patients with idiopathic generalized epilepsy, and could identify that the former group display distributed patterns of cortical E/I imbalances, together with reductions in subcortical drive^18^. Despite limited data on the interaction between these chronic alterations in intrinsic brain function and transient epileptic events, it is plausible that a chronic functional imbalance, as indicated by reduced INT findings, may also ultimately predispose brain networks to exhibit transient epileptic states^51^. An earlier study reported state-dependent changes in global network dynamics, as indicated by a decrease of neuronal synchronization during seizures and increases prior to seizure termination^6^, consistent with the above conjecture.

Structural and diffusion MRI studies in TLE have previously mapped patterns of cortical and subcortical morphological changes as well as atypical fiber diffusivity in deep and superficial white matter. Capitalizing on an advanced multimodal MRI processing and feature fusion framework^22^, our work examined associations between INT changes and MRI-derived group differences in grey matter morphology and white matter diffusivity in the same participants. Patterns of grey matter atrophy and white matter diffusion alterations observed in the current work were largely in line with prior studies, suggesting bilateral and widespread cortico-subcortical atrophy, and a predominance of temporo-limbic alterations in white matter diffusivity and microstructure^10, 12, 18, 19, 25, 26^. Notably, while the spatial patterns of INT reductions in TLE were similar after correcting for both grey and white matter changes in the same regions, effect sizes of TLE-related INT changes decreased, particularly in subcortical regions and in the hippocampus when correction was applied. Such findings overall demonstrate that INT changes provide unique information that is not comprehensively explained by grey matter morphology and proximal white matter microstructure, but that there nonetheless exists shared variance between structural and functional alterations. Marked modulations of functional matrices by structural features, as observed in the mesiotemporal lobe, are in line with the role of the hippocampus as the pathological core of TLE, and its position as a convergence zone of distributed functional processing streams^52, 53^. As such, these findings support prior work in TLE provide a system-level functional complement to previous work showing marked modulatory effects of hippocampal pathology on large-scale white matter network topology and connectome-informed network communication models^10, 53^.

Performing an age by group interaction analysis provided a cross-sectional window into progressive functional changes in TLE, and their differences from ageing effects in controls^12^. Specifically, we found pronounced age by group interactions on INT in bilateral primary cortices and ipsilateral temporal regions, with more marked negative effects of age in TLE than in controls. Patterns in patients were also significantly associated with disease duration, which may indicate multiple parallel processes, notably the cumulative effects of ongoing seizures. Despite an incompletely understood association between seizures and network function, seizure-related reorganization of brain networks have been supported by experimental studies in animal models of limbic epilepsy^54^. Moreover, MR spectroscopic work in patients with TLE suggested associations between seizure burden and measures of neural integrity^55^, and there is relatively convincing evidence from cross-sectional and longitudinal analyses of grey matter morphology indicating progressive atrophy in TLE patients ^56, 57^. On the other hand, except for prior research showing progressive effects on *in vivo* PET-derived glucose metabolism data^58^, findings demonstrating age-related effects on intrinsic function at the whole-brain level in TLE have remained scarce until now, and our study thus provides a potential region-specific functional complement to the extant structural literature. Arguably, and beyond seizure related effects, our findings showing more marked INT reductions in patients with advanced age and duration may also signify effects of continued treatment with anti-seizure medication, or result from overall challenges to quality of life and occupational independence that TLE patients may face^51^. Several studies have shown associations between treatment with anti-seizure medication and measures of brain structure^59^ as well as function^51^. However, a precise isolation of specific drug-effects remains difficult given that patients may often be on polypharmacotherapy and/or switching drug combinations throughout their clinical course. Accelerated ageing processes in measures of intrinsic brain function could furthermore be in line with observations of vascular alterations in TLE^60^ and emerging histopathological reports suggesting a higher presence of neurodegenerative deposits, including misfolded tau protein, in surgical tissue resected from TLE patients^61^. Future work tracking changes in brain function and structure longitudinally, alongside careful monitoring of clinical and cognitive phenotypes will help to further sharpen our understanding of causes and consequences of epilepsy-associated progression. To account for a likely broad range of susceptibility and resilience factors across patients, such efforts would certainly benefit from multi-site data aggregation and dissemination initiatives such ENIGMA-Epilepsy^62^.

Leveraging supervised machine learning, we demonstrated that classifiers informed by INT could distinguish TLE from controls and lateralize the seizure focus with high accuracy. These findings complement earlier efforts in epilepsy classification and focus lateralization using a wide range of features from structural MRI, diffusion MRI, and rs-fMRI connectivity analysis^63-65^. Notably, we specifically assessed cross-site prediction and confirmed adequate generalization, contrasting most of the prior that evaluated functional classifiers within single sites. Informative features included mesiotemporal areas but also spanned beyond these, likely reflecting the known system-level involvement of TLE^4, 12^. Collectively, these findings highlight a promising utility of intrinsic time scale parametrization for personalized diagnostics in TLE, and motivate future biomarker discovery efforts that combine different structural and connectional measures with biologically meaningful indices of brain function, such as INT.

## Glossary

ACF: autocorrelation function
AUC: area under the curve
BACC: balanced accuracy
DWI: diffusion weighted imaging
EEG: electroencephalography
FA: fractional anisotropy
FD: framewise displacement
ILAE: International League Against Epilepsy
INT: intrinsic neural timescales
MD: mean diffusivity
ROC: receiver operating characteristic
rs-fMRI: resting-state functional MRI
TLE: temporal lobe epilepsy.

## Supplementary results

**Figure S1.**
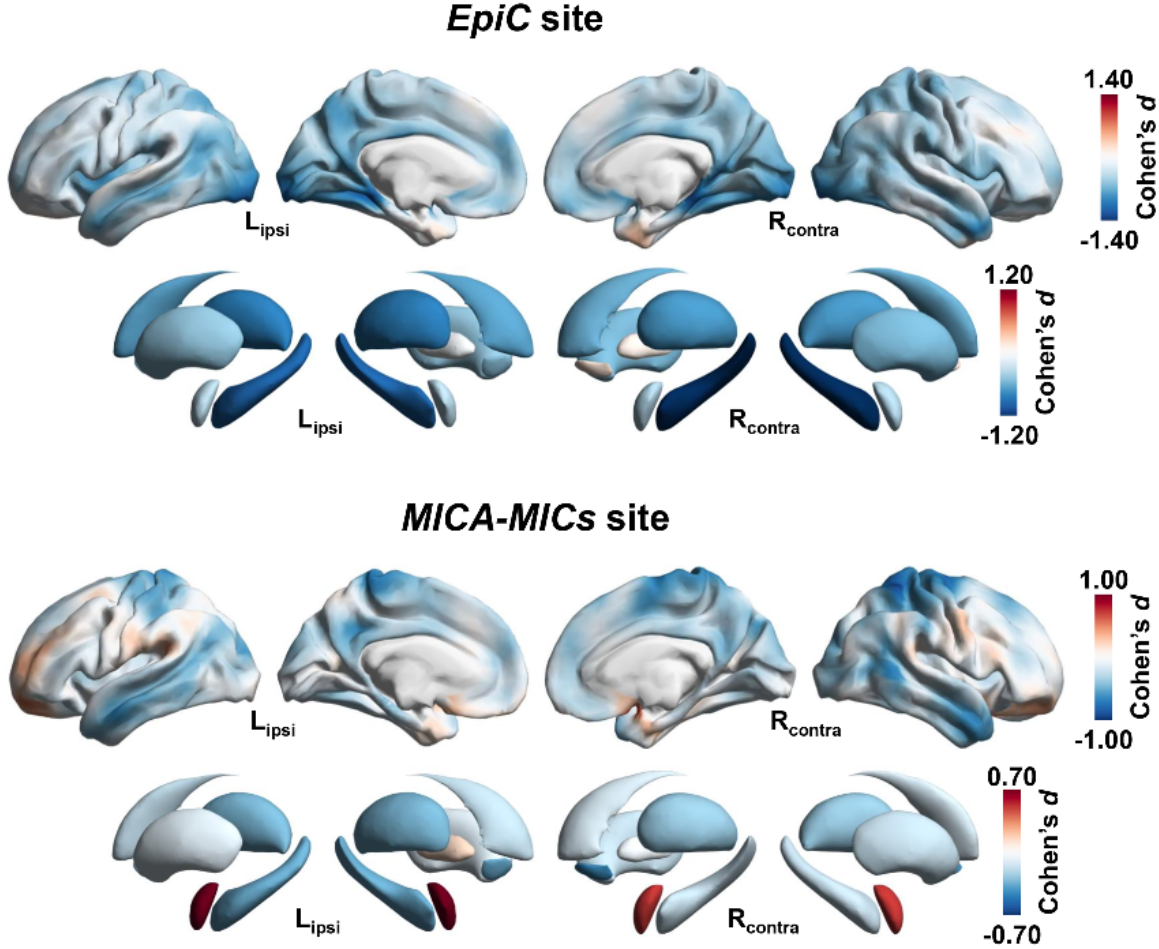
Between-group differences in regional intrinsic neural timescales (INT) carried out separately in the *EpiC* and *MICA-MICs* site, after correcting for age and sex.

**Figure S2.**
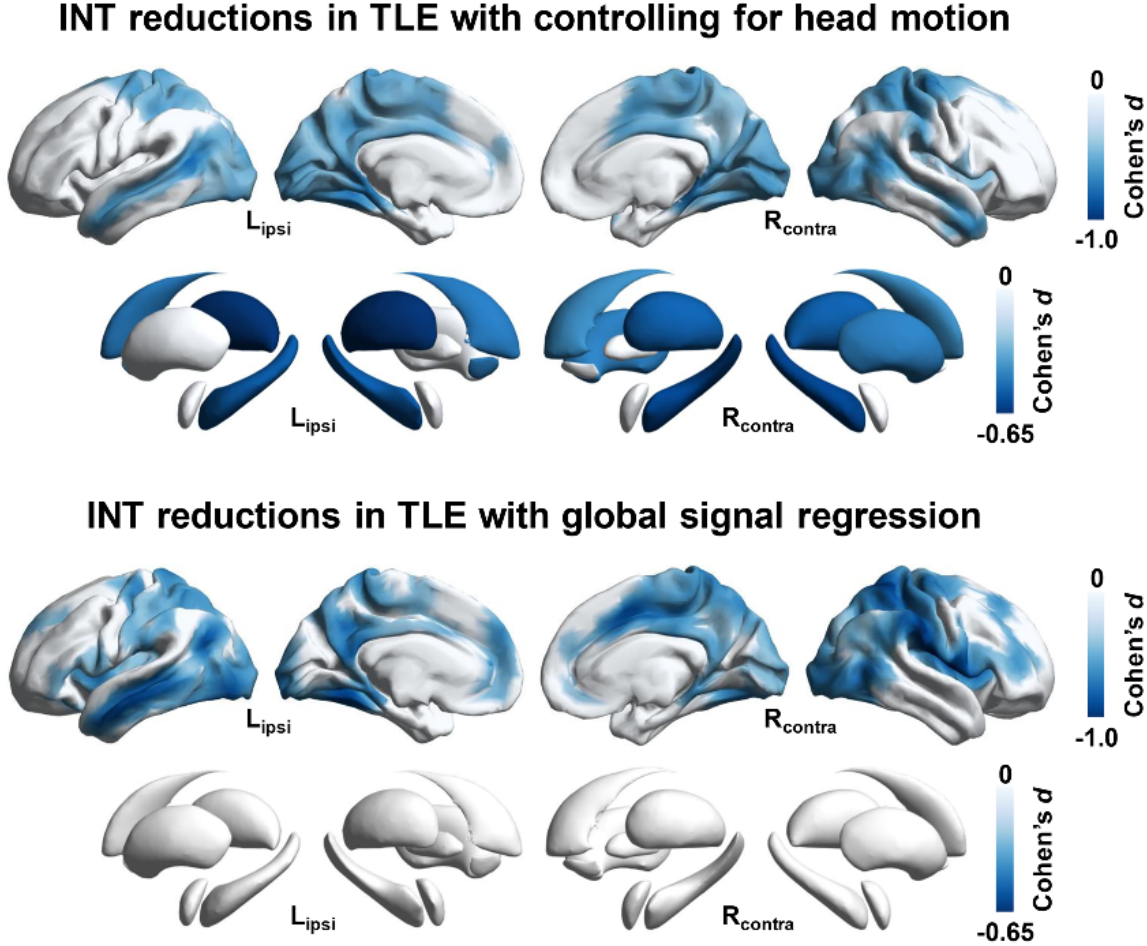
Between-group differences in regional intrinsic neural timescales (INT) after additionally regressing out head motion (*top*) or the global mean signal (*bottom*). Findings have been corrected for multiple comparisons at a family-wise error (FWE) rate < 0.05 for cortices, and at a false discovery rate (FDR) < 0.05 for subcortices and the hippocampus.

**Figure S3.**
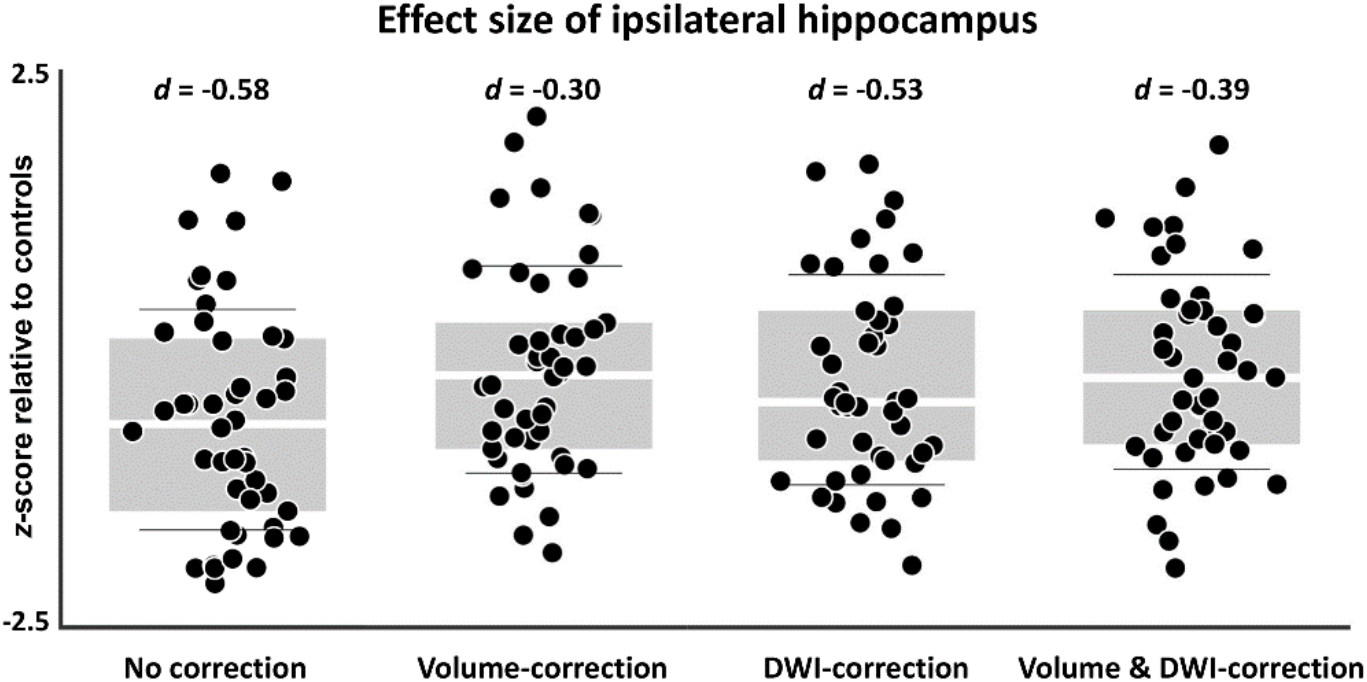
Effect sizes of intrinsic neural timescales (INT) reductions of the ipsilateral hippocampus in TLE with/without controlling for grey matter morphological and white matter microstructural alternations.

**Figure S4.**
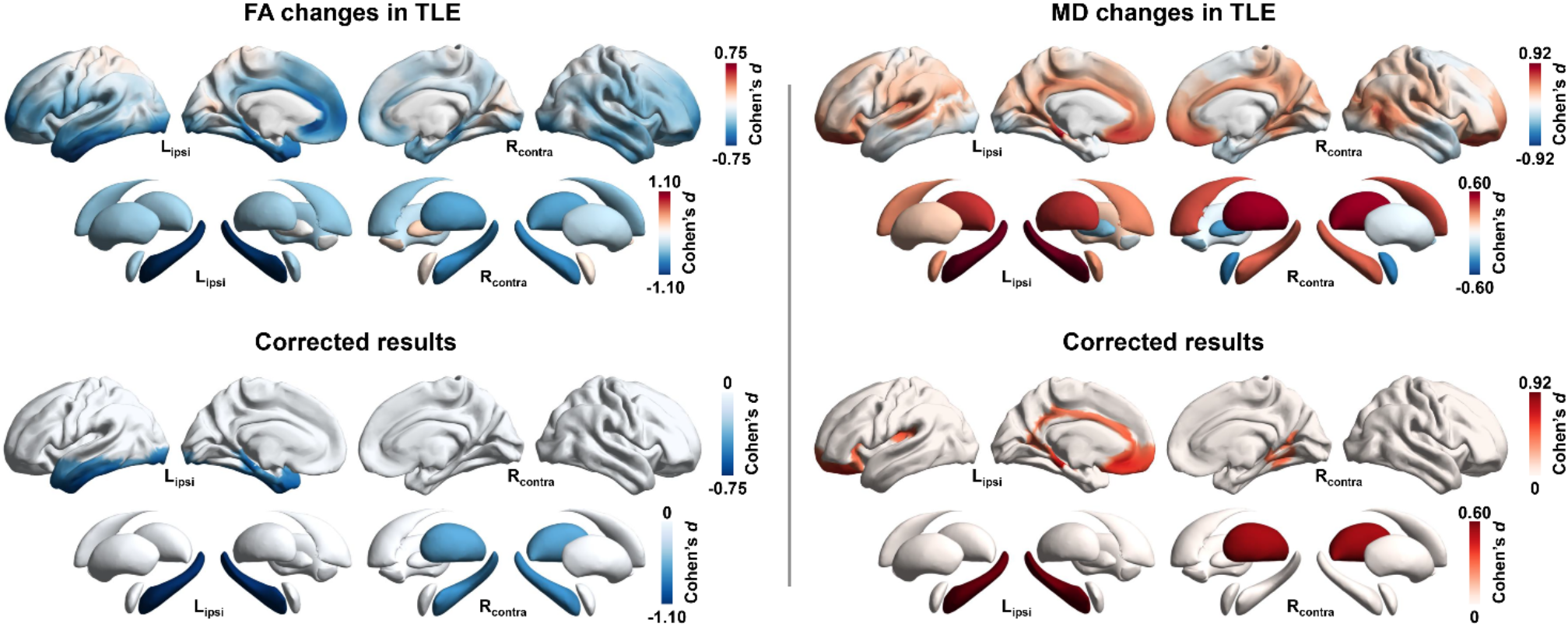
Patterns of univariate superficial white matter microstructural perturbations in TLE. Compared to healthy controls, TLE patients presented with markedly decreased fractional anisotropy (FA) and increased mean diffusivity (MD), with dominant effects ipsilateral to the focus. To correct for multiple comparisons, cortical regions were thresholded at a family-wise error (FWE) rate < 0.05, and subcortical regions and the hippocampus were thresholded at a false discovery rate (FDR) < 0.05.

**Figure S5.**
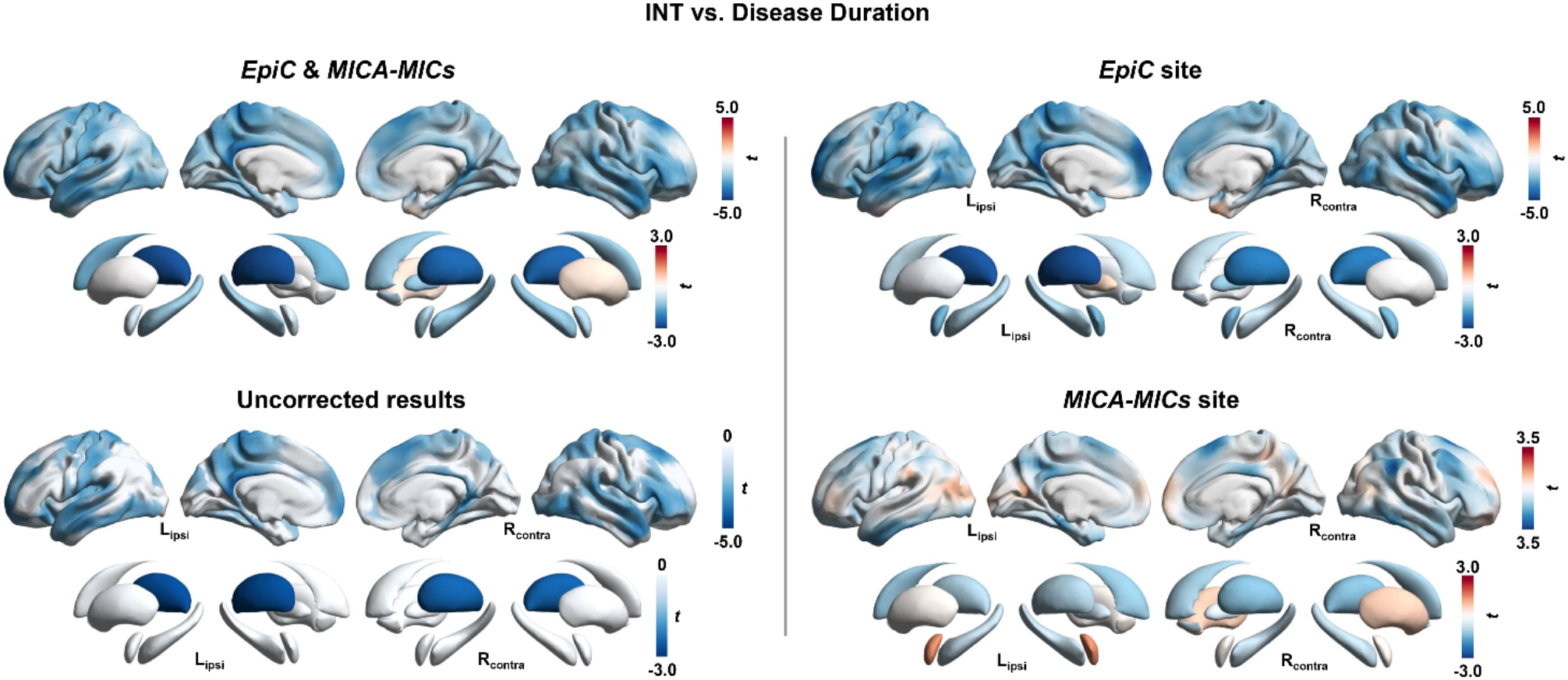
Main effects of epilepsy duration on intrinsic neural timescales (INT) in TLE following multiple comparisons corrections at a family-wise error (FWE) rate < 0.05 for cortical regions, and at a false discovery rate (FDR) < 0.075 for subcortical regions and the hippocampus.

**Figure S6.**
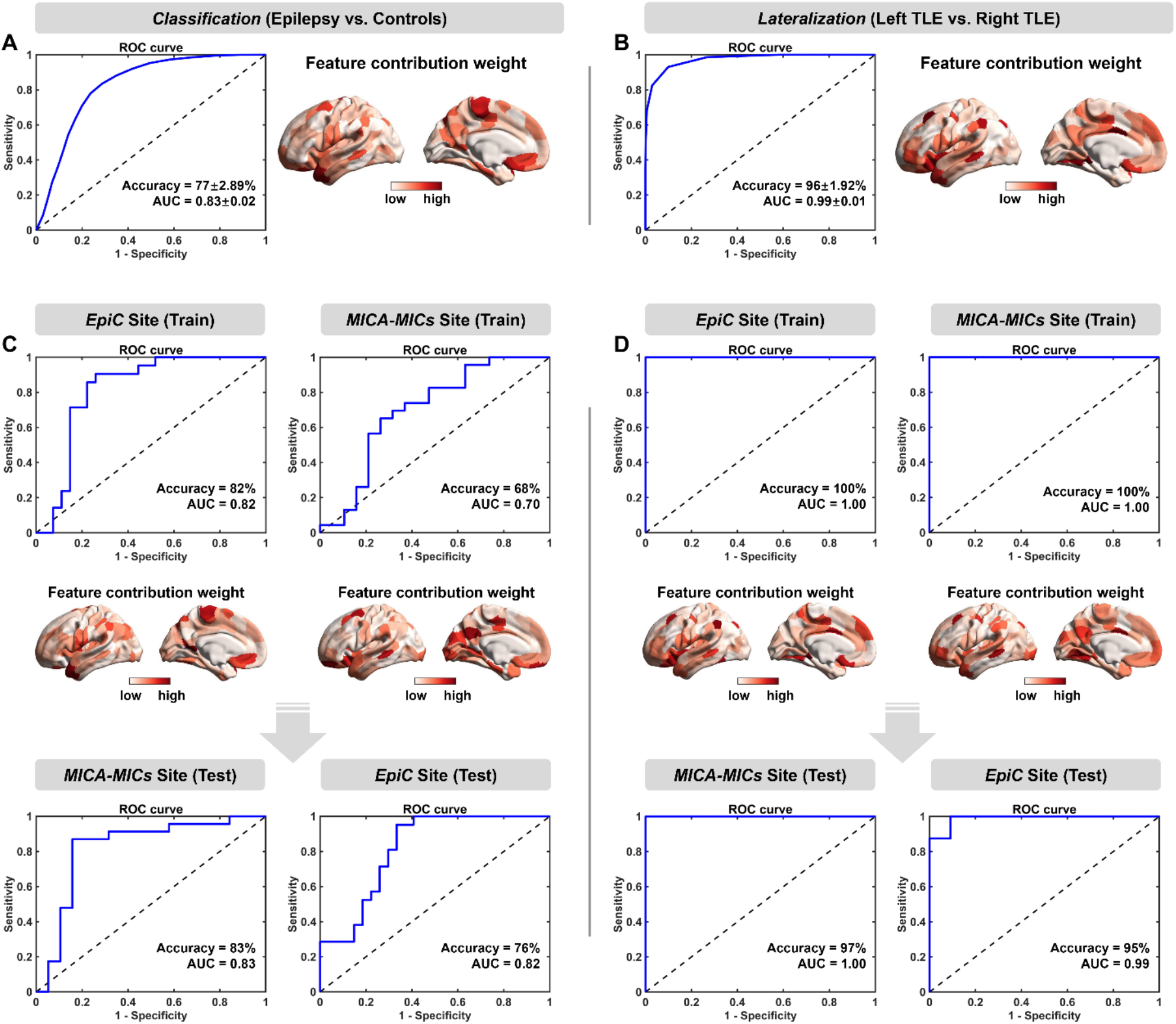
Classification and lateralization analyses using cortical intrinsic neural timescales (INT). Leveraging support vector machine (SVM) algorithm to build models for TLE-*vs*-controls classification (left panel), and models for left-*vs*-right TLE lateralization (right panel). **(A)** and **(B)** Performance of classification and lateralization models trained and tested across two datasets, using five-fold cross-validation. Average receiver operating characteristic (ROC) curves, normalized feature contribution weight maps, balanced accuracy, and area under the curve (AUC) (mean ± SD) across 100 iterations. **(C)** and **(D)** Performance of classification and lateralization models that were trained on one dataset and tested on the other.

**Table S1.** Classification and lateralization performance using cortical features.

|  | *EpiC & MICs* |  | *EpiC (Train) → MICs (Test)* |  |  |  | *MICs (Train) → EpiC (Test)* |  |  |  |
| --- | --- | --- | --- | --- | --- | --- | --- | --- | --- | --- |
|  | BACC <sup>a</sup> | AUC <sup>a</sup> | BACC | AUC | BACC | AUC | BACC | AUC | BACC | AUC |
| Classification | 77±2.89% | 0.83±0.02 | 82% | 0.82 | 83% | 0.83 | 68% | 0.70 | 76% | 0.82 |
| Lateralization | 96±1.92% | 0.99±0.01 | 100% | 1.00 | 97% | 1.00 | 100% | 1.00 | 95% | 0.99 |

Abbreviations: BACC = balanced accuracy; AUC = area under curve.
<sup>a</sup> BACC and AUC in mean ± SD.

